# Pollinator and flower morphology interact to influence pollen receipt

**DOI:** 10.1101/2025.05.16.654420

**Authors:** Ethan Newman, Allan Ellis, Bruce Anderson

## Abstract

Pollinators are important drivers of floral divergence and speciation in plants. Here we investigate the effects of floral trait variation and trait matching on pollen receipt in two different plant species, *Tritoniopsis revoluta* (Iridaceae) and *Nerine humilis* (Amaryllidaceae) each visited by long-proboscid nemestrinid flies and solitary bees across their ranges. Using single visits by pollinators to flowers in populations with experimentally increased variance in floral morphology, we demonstrate that pollinators with different morphology affect pollen receipt differently through their interactions with floral traits. This study provides details of how the mechanical fit between floral and pollinator morphology may drive floral divergence through its effects on pollen deposition, both within populations using multiple pollinators and between populations with contrasting pollinators.

## Introduction

Pollinator-mediated selection plays an important role in driving floral divergence (Van der Niet et al. 2014; Johnson 2025). In support, several studies have demonstrated local adaptation and consequent divergence of geographic floral forms pollinated by functionally different pollinators (Robertson and Wyatt 1990; Sun et al. 2014; Newman et al. 2015; Johnson et al. 2025). As effective pollination requires precise placement of pollen on a part of the pollinator’s body that will contact stigmas of subsequently visited flowers, divergence often involves traits that determine the mechanical fit of flowers and functionally different pollinators.

We studied *Nerine humilis* (Jacq.) (Amaryllidaceae) and *Tritoniopsis revoluta* (Burm.f.) (Iridaceae), both of which have divergent floral ecotypes associated with different pollinators. We chose these species because they appeared to adapt to variation in pollinator morphology by utilizing very different kinds of floral morphology (i.e., style lengths versus floral tube length). Both species are pollinated by nectar foraging nemestrinid flies with long proboscides, but also by bees which have much shorter proboscides, and exhibit geographic variation in floral traits associated with the morphological fit between flowers and their pollinators (Newman et al. 2014). In *N. humilis*, pollen transfer occurs on the abdomen of insects (Fig. 1a, b), and effective pollination is achieved when the functional pollinator length matches anther and style length (Newman et al. 2014). As nectar access is not restricted by a long floral tube, it is accessible to pollinators regardless of their proboscis length (Newman et al. 2015). In contrast, *T. revoluta*, like many other taxa pollinated by long-proboscid insects, has evolved long nectar tubes. This would force insects to enter the floral chamber upon foraging (Fig. 2), where the stigma is able to receive pollen from the thorax of the fly (Anderson and Johnson 2008; Muchhala and Thomson 2009; Pauw et al. 2009). Consequently, selection for tighter mechanical fit should favor *T. revoluta* individuals with nectar tubes that are equal to, or longer than the proboscides of pollinators.

**Figure 1.**
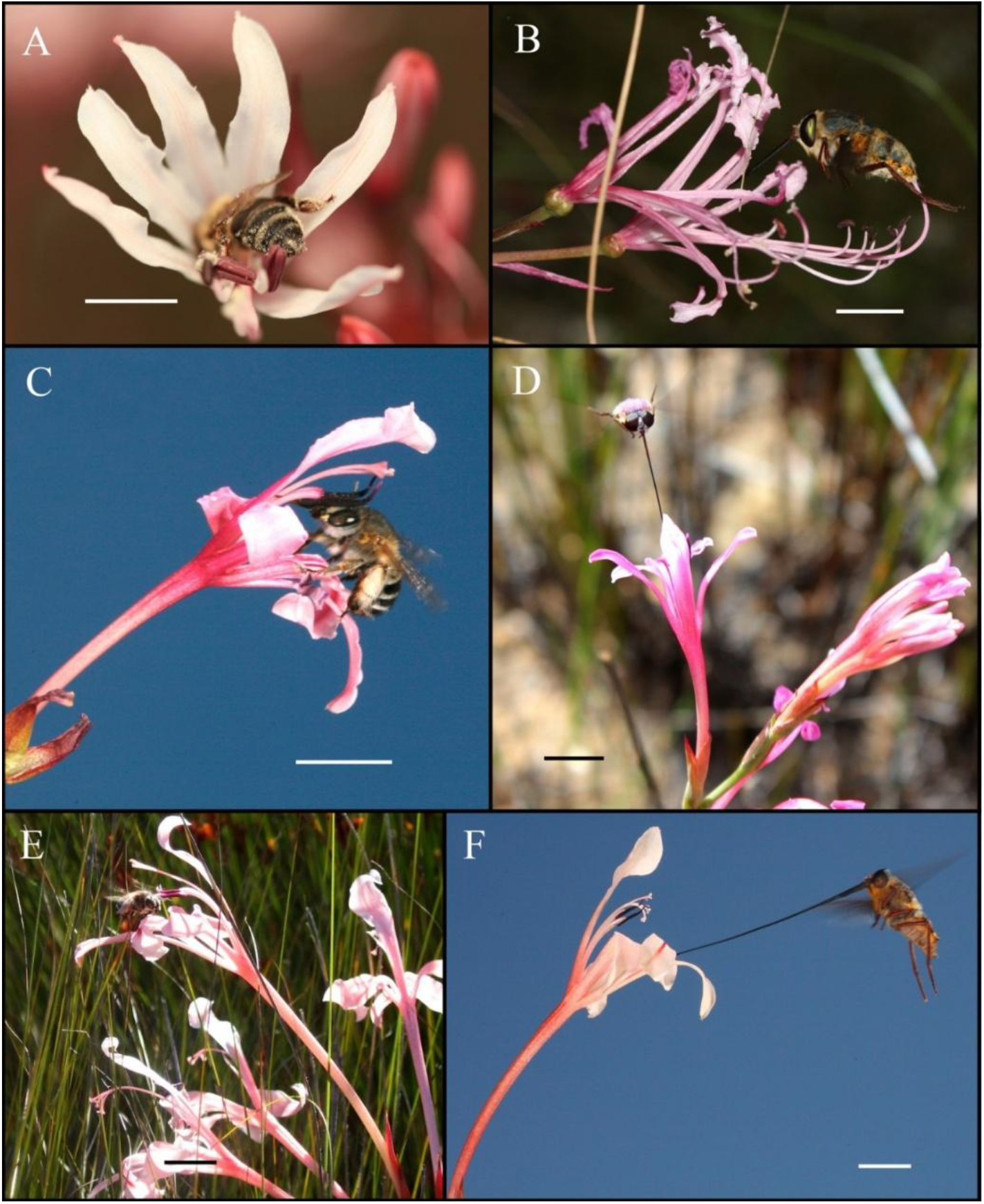
Pollinators found in different populations of *Nerine humilis* and *Tritoniopsis revoluta*. (a) Honeybee pollinators were the sole pollinators of short-styled *Nerine humilis* plants from Swellendam. (b) The long proboscid fly *Prosoeca longipennis* was the dominant pollinator visiting long-styled *N. humilis* from the Baviaanskloof. *Amegilla* bees (c) and an unidentified *Prosoeca* species (d) were common visitors to *Tritoniopsis revoluta* flowers with short tubes in the Swartberg. *Amegilla* bees (e) and the long proboscid fly *P. longipennis* were the two most frequent visitors to very long tubed *T. revoluta* flowers from Barrydale. Scale bar = 1cm. Photos by E. Newman and B. Anderson, except (d) by Theo Busschau.

**Figure 2.**
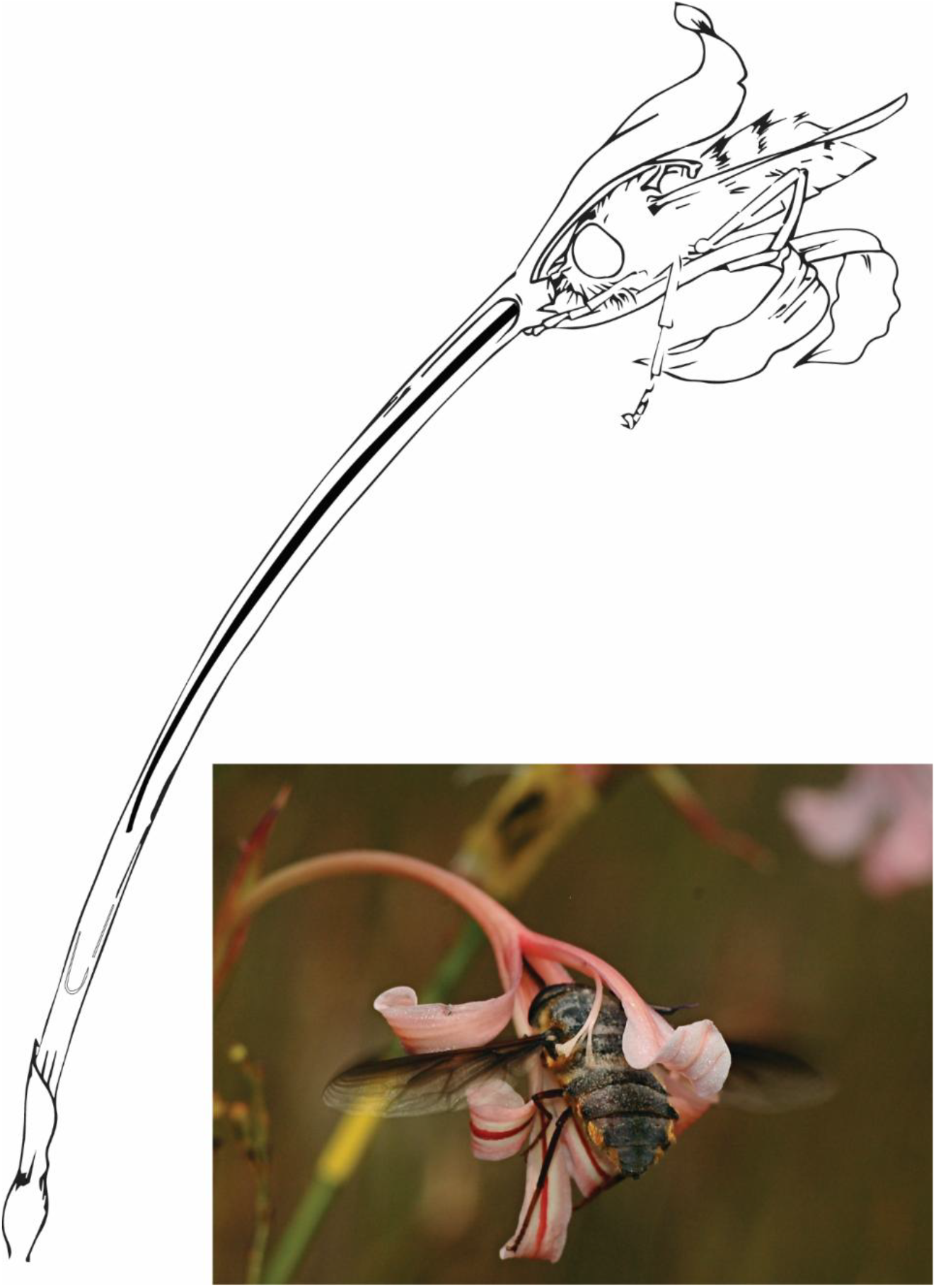
Illustration and photographic insert (photo by Caitlin von Witt) demonstrating the fit between a *Tritoniopsis revoluta* flower and *Prosoeca longipennis*, a long proboscid fly pollinator at Barrydale. Only when a visitor enters the floral chamber created by the tepals, does its body make contact with the receptive stigma which resides within the chamber.

By introducing flowers from morphologically diverged populations, we were able to experimentally increase the phenotypic variation available for pollinator morphology to interact with. This helps circumvent a frequent problem encountered by many phenotypic selection studies: strong selection often eliminates much of the phenotypic variation in populations, making it hard to detect selection (Waterman et al. 2023). By introducing low numbers of extreme phenotypes to existing population variation, we mimic the process by which extreme traits may arise through a rare mutation or migration (Trunschke et al. 2020). We then quantified pollen deposition after single visits by different pollinators to newly opened flowers that differed in morphology. We repeated this across multiple localities with divergent floral variation and pollinator communities to compare performance surfaces associated with differing levels of plant-pollinator trait matching.

Our primary objective with this experiment was to provide much needed insights into how the functional relationships between flower and pollinator morphology affects pollinator-mediated selection, both within populations, but also between populations. Here we investigate how variation in *N. humilis* style length and *T. revoluta* tube length affects pollen receipt when visited by pollinators with different morphology. The two plant taxa, act as replicates to test the hypothesis that pollinators contrasting in their functional fit with flowers, should generate divergent patterns of pollen deposition (different pollen performance landscapes), see Fig. 3 for specific predictions.

**Figure 3.**
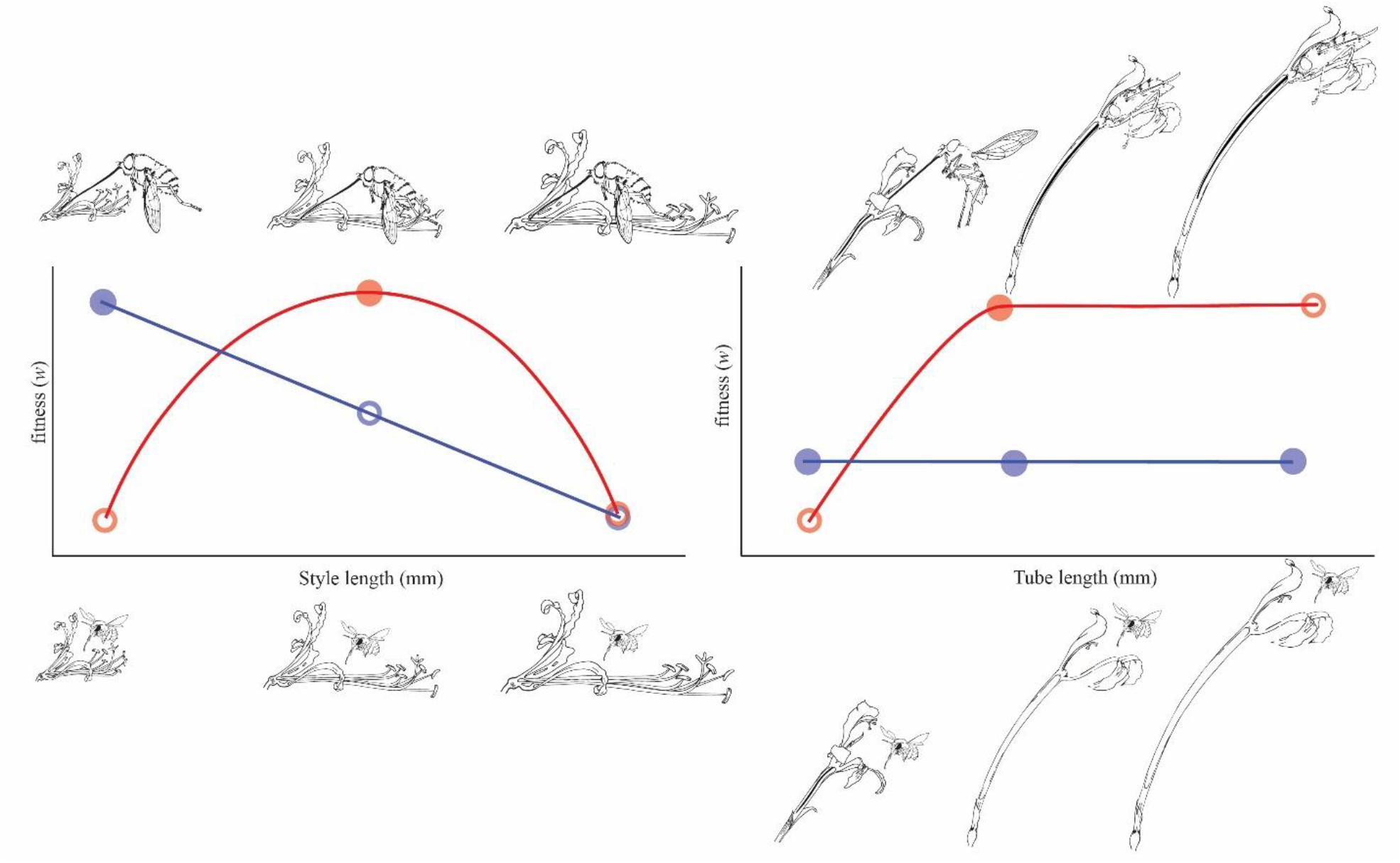
Hypothetical differences in the shapes of selection surfaces imposed by long proboscid flys’ (red) and short proboscid insects (blue). In *Nerine humilis* (A), we expect visits by long proboscid fly pollinators (top illustrations) to produce a stabilizing selection signature. Here we expect plants with intermediate styles to make good contact with the abdomen of the fly and as a result experience a higher fitness advantage over flowers with short and long styles. In the presence of short proboscid insect visitors (bottom illustrations) we expect a negative directional selection signature where extreme short style flowers experience a higher fitness advantage over intermediate and long style flowers. In *Tritoniopsis revoluta* (B), visits by long proboscid flies are expected to lead to a saturating curve associated with directional selection. This is because pollinator body position relative to floral reproductive structures does not change once tube length exceeds pollinator tongue length. However, we expect no selection on *Tritoniopsis revoluta* tube length after visits by bees, as all bees are required to enter the gullet of *T. revoluta* to obtain the nectar which wells up within the tubes. Dotted lines for both *Nerine humilis* and *Tritoniopsis revoluta* indicate style/tube lengths where the mechanical fit between flowers and pollinators is optimized.

## Materials and Methods

### Study species and experimental localities

*Nerine humilis* (Amaryllidaceae) and *T. revoluta* (Iridaceae) are protandrous geophytes (Ros et al. 2011; Newman et al. 2015) that flower from March-June in the winter rainfall region of South Africa (Fig. 1). *N. humilis* was studied in Swellendam (−34.184574°S, 20.291919°E), where flowers had short styles (mean ± SD = 26.8±4.17mm) and were visited almost exclusively by nectar foraging honeybees (*Apis mellifera*) with short proboscides (<5mm) (Newman et al 2015), and at Nuwekloof pass in the Baviaanskloof Megareserve (−33.511291°S, 23.642786°E), where it has much longer styles (mean ± SD = 42.03±6.15mm) and is primarily visited by *Prosoeca longipennis and P. ganglebauri* flies with proboscides in excess of 20mm (Newman et al. 2015). These two populations differ statistically from one another in style length, the functional trait associated with mechanical fit (*t*=-18.17, *df*=149, *P*<0.001).

*T. revoluta* was studied at two populations differing in tube length (*t*=-17.18, *df*=135, *P*<0.001) and pollinator community composition. Plants from the Swartberg mountains (−33.362594°S, 22.068132°E) had short tubes (mean ± SD = 25.20±6.81mm) and are visited by solitary *Amegilla* bees with short proboscides (<12mm) and a long proboscid fly species (*Prosoeca* sp.) with a longer proboscis (>15mm). Plants from Barrydale (−33.935997°S, 20.679453°E) had much longer tubes (mean ± SD = 66.38±7.33mm) and are visited by solitary *Amegilla* bees and by a fly, *P. longipennis*, with an exceptionally long proboscis which can exceed 60mm in length. In both populations, the proboscis lengths of the flies are closely matched with the tube lengths of the flowers (Anderson et al. 2014) and it is expected that these flies are able to access most of the nectar found within the corolla tubes. However, the solitary bees have proboscides many times shorter than the *T. revoluta* tubes and appear to obtain small amounts of nectar when the nectar accumulates in the tubes and rises to within their reach (de Merxem et al. 2009).

### Field experiments and floral trait measurements

To study the effects of pollinator and floral morphology on pollen deposition, we added floral variation to the experimental localities by translocating individuals from other populations. Imported plants were targeted to span the range of floral trait variation found within each species so that we could construct performance surfaces that encompassed broad ranges of floral variation. Virgin flowers were then exposed to a single floral visitor after which the stigma was harvested and the deposited pollen counted.

For *N. humilis* at Swellendam, we supplemented 54 local experimental plants (mean style length =26.34mm, range = 20.18-36.35mm) with 50 short-styled flowers from Rooibrug, -34.067066°S, 20.390489°E, (mean style length =24.84mm, range =18.36 -30.93mm) and 41 long-styled flowers from Suurbraak, -34.015907°S, 20.598108°E, (mean style length =38.35mm, range = 30.93-46.99mm), giving a total of 145 experimental flowers. At the Nuwekloof Pass, Baviaanskloof, 32 local inflorescences (mean style length = 39.65mm, range = 21.92-51.52mm, n=32) were supplemented with 37 short-styled flowers from Napier (mean style length = 28.14mm, range = 20.07-41.56mm), giving a total of 69 experimental flowers.

For *T. revoluta* at Barrydale, we used 21 local long-tubed experimental flowers (mean tube length = 70.00mm, range = 62-80mm) together with longer tubed flowers from Tradouws 1, -33.946883°S, 20.698283°E (mean = 85.75mm, range = 82-91mm, n= 4), and shorter tubed flowers from Gysmanshoek long, -33.930278°S, 21.0765°E (mean = 51.67mm, range = 40-60mm, n = 6) and Gysmanshoek short, -33.9325°S, 21.071944°E (mean = 36.22, range = 29-47mm, n= 23). Similarly, at Swartberg, with shorter tubes on average, we used 20 native flowers (mean tube length = 21.25, range = 14-26mm), supplemented with flowers from Barrydale (mean = 67.00mm, range = 61-79mm, n = 6), Gysmanshoek long (mean = 54.25mm, range = 49-60mm, n = 12), Gysmanshoek short (mean = 34.16, range = 27-47, n= 46) and Tradouws 2, - 33.98715°S, 20.722183°E (mean = 10.93mm, range = 6-13mm, n = 15). This gave us a total of 54 flowers at Barrydale and 99 flowers in the Swartberg. The introduced plants represented less than 5% of the total plants in the population.

Inflorescences used for single visits were cut so that they contained buds at the same stage of development (see replication above). All experimental inflorescences (including local ones) were allowed to develop in water for 2-3 days until the flowers opened and stigmas became receptive. Flowers were emasculated when the anthers matured to prevent possible contamination of local gene pools with translocated pollen but also to prevent pollen movement within a flower. Once stigmas were receptive, inflorescences, containing a single experimental flower each, were placed in test tubes filled with water and mounted on skewer sticks.

Experiments ran between three and five days at each locality, between 0900hrs and 1200hrs when pollinators were most active. After a visit to an experimental flower, the visited flower was immediately covered with netting to prevent further visits. After each day, the number of conspecific pollen grains on the stigmas of visited flowers was counted under a dissection microscope. This was done using fuchsin gel for *Tritoniopsis*, but the large pollen grains of *Nerine* were counted without the use of fuchsin gel (as in Newman et al. (2015)).

Style lengths of visited *N. humilis* flowers were measured in female phase from the top of the ovary in the nectar chamber to the tip of the stigma (see Newman *et al*., 2015) and for *T. revoluta*, tube length was measured from the top of the ovary to where the corolla flares (see Anderson *et al*., 2014). We also measured the relevant functional traits of pollinators euthanized using Potassium cyanide fumes, following their capture with an insect net. For insects visiting *T. revoluta*, we measured proboscis length (i.e. the distance from the base to the tip of the fully extended proboscis) as the mechanical fit trait (Anderson and Johnson, 2008). Because *N. humilis* places pollen on the end of the pollinator’s abdomen, we measured the functional body length of insects, (proboscis length + body length) accounting for the angle at which insects hold their proboscis whilst foraging (see Newman *et al*., 2015). All traits were measured to the nearest millimeter using a set of digital calipers.

### Statistical analysis

Performance surfaces are visualizations of the effects of phenotypic trait variation on components of fitness (Arnold 2003), and have long-been used to understand the mechanisms behind microevolutionary divergence on important functional traits within various systems (Wade and Kalisz 1990). In this study, we use performance surfaces to isolate the effects of different pollinators on pollen receipt across a range of possible floral phenotypes.

By estimating pollen receipt and relating it to individual floral trait values, we explored how two distinct functional pollinator groups, i.e. long-proboscid flies (*Prosoeca*) and short-proboscid bees influence pollen deposition in two divergent populations of two different plant species. We modelled pollen receipt surfaces for the two *N. humilis* populations using thin-plate spline generalized additive models (Schluter and Nychka 1994; Arnold 2003), using the “gam” command, implemented in the package mgcv (Wood 2017). As overdispersion and non-normality of residuals in Poisson distribution models were discovered using the “DHARMa” package (Hartig and Hartig 2017), we used negative binomial error distributions with log link functions in our models, which showed no evidence of overdispersion or non-normality after model diagnostics were reassessed. In these models, Lambda, which controls the “wiggliness” of the curve, was penalized using REML (Wood 2017).

For the two *T. revoluta* populations, each visited by two different functional pollinators within the same experimental population (bees and long proboscid flies), we used the same analysis and approach as for *N. humilis*, except that separate models were run for each site. First, we ran a global model that included pollen deposition from all pollinators. Next, to determine whether pollinator functional types (bees versus flies) contribute to statistically different pollen deposition surfaces, we ran a model treating pollinator functional type as a “fixed effect” using the “by” command. Statistical significance between smooths generated by different pollinators was determined using the “anova.gam” command (Wood 2013).

We further obtained directional (*β*) and quadratic (*γii*) performance coefficients from generalized linear models (GLM) that included pollen deposition as the response and mean standardized style length for *N. humilis* and tube length in *T. revoluta* as predictor variables, using the approach of Morrissey and Goudie (2022). These models used a negative binomial error distribution with a log-link function. We used this approach over that of Lande and Arnold (1983), due to overdispersion issues in our initial models, which was discovered using the DHARMa package (Hartig and Hartig 2017). GLM’s were implemented in the “MASS” package (Ripley et al. 2013), and all analyses were done in R (Core Team, 2024).

## Results

For *Nerine humilis* in Swellendam, we recorded 145 single visits – all by honeybees (mean functional body length±SE = 15.70±0.28 mm, n=14). In contrast, 69 single visits were made to *N. humilis* by two functionally similar long-proboscid flies at Nuwekloof Pass (43 by *P. ganglebauri:* mean functional body length±SE = 47.60±5.57mm, n=3 and 26 by *P. longipennis:* mean functional body length± SE = 39.72±1.24mm, n=10).

For *T. revoluta* in the Swartberg, 99 single visits were received, with 42 visits made by bees (14 *Amegilla* sp. and 28 *Lasioglossum* sp.), all with proboscis lengths shorter than 12mm (mean proboscis length± SE =11.20±0.30mm, n=8). The rest of the visits (n = 57) were made by a long proboscid fly, *Prosoeca* sp. (mean proboscis length± SE = 23.36±1.12mm, n=11). *Tritoniopsis revoluta* in Barrydale received 54 single visits: 32 by the solitary bee, *Amegilla* sp. (mean proboscis length±SE = 11.50±0.61mm, n=12) and 22 visits by the long-proboscid fly *P. longipennis* (mean proboscis length±SE = 68.9±0.75mm, n=6).

### The effects of floral and pollinator morphology on pollen deposition in Nerine humilis

At the Swellendam population, where *N. humilis* is visited by honeybees, the thin plate spline suggests that pollen receipt decreased with increasing style length (χ^2^ =14.72, edf =1, p < 0.001, Fig. 4A). This decreasing trend was also evident when the data were analyzed using a univariate generalized linear model (Table 1), suggesting negative directional selection on style length mediated by bees.

**Table 1.**
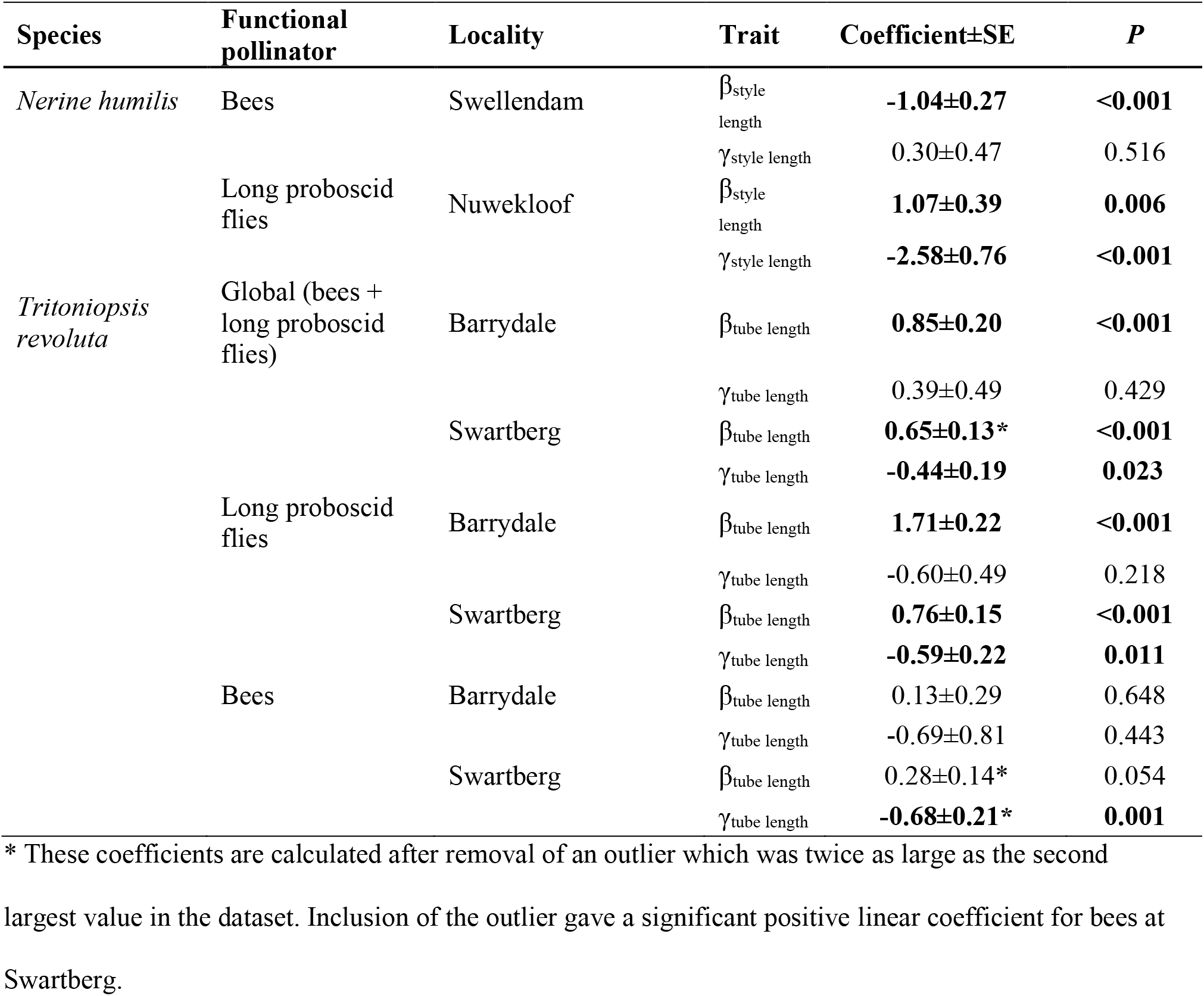
Standardized directional (*β*) and quadratic (γ) pollen receipt coefficients on style length in *Nerine humilis* and on tube length in *Tritoniopsis revoluta*. Pollen receipt differentials were estimated using generalized linear models with absolute pollen receipt as the response and standardized tube or style length as the explanatory variable. Estimates highlighted in bold are significant when *P*<0.05.

**Figure 4.**
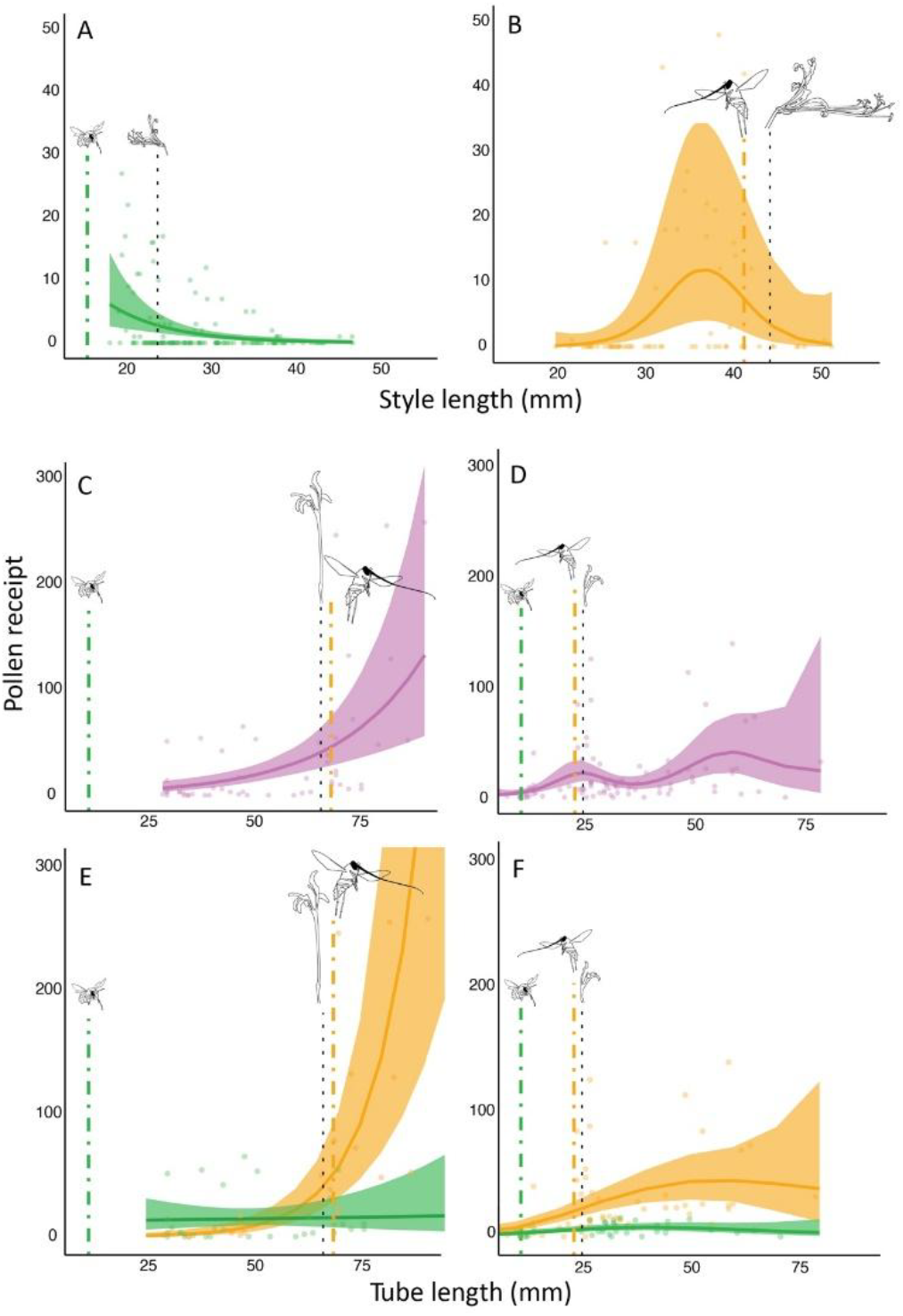
Generalized additive model (GAM)-based pollen receipt surfaces mediated by honeybees on *Nerine humilis* style length at Swellendam (a) and long-proboscid flies, *Prosoeca longipennis* at Nuwekloof pass (b). *Tritoniopsis revoluta* populations are each visited by functionally distinct pollinators, instead of one, that differ in pollen deposition performance along the axis of experimental trait variation. Hence, panel (c) represents a global pollen receipt surface that combines pollen deposition from both short and long proboscid pollinators from Barrydale, and panel (d) represents the global pollen receipt surface from the Swartberg. Panel (e) represents separate pollen receipt surfaces for *Amegilla* bees and long proboscid flies at Barrydale, whilst (f) represents surfaces for solitary bees and flies with medium proboscis lengths at Swartberg. The solid line represents the cubic spline and the shaded outline, 95% confidence intervals. For each locality, population mean style/tube length is represented by the dotted black line, and mean functional pollinator length with either green (bees) or orange dashed lines (long proboscid flies) (*see* results for measurements).

At Nuwekloof pass, where *N. humilis* is visited by long proboscid flies, the thin plate spline shows a significant non-linear relationship between pollen received and style length (χ^2^ =12.49, edf =2.58, *P* = 0.008, Fig. 4B), with an optimum around 35mm that is slightly shorter than the average functional lengths of the visiting flies at this site (Fig. 4B). The clear optimum suggests stabilizing selection (Mitchell-Olds and Shaw 1987) where intermediate styles receive more pollen, than either short or long-style plants (see predictions, Fig. 3), also supported by a significant negative quadratic coefficient (Table 1). Furthermore, there was also a significant positive linear coefficients indicative of positive directional selection on style length, suggesting a general increase of pollen grains received with increasing style length (Table 1).

### The effects of floral and pollinator morphology on pollen deposition in Tritoniopsis revoluta

Global pollinator receipt surfaces, where single visit data from all pollinators were combined at a site, showed significant positive trends at both *T. revoluta* sites (Barrydale: χ^2^ =19.53., edf =1.00, p <0.001; Swartberg: χ^2^ =38.68, edf =5.02, *P* <0.001, Fig. 4C and D). These positive trends were supported by significant positive GLM linear coefficients at both Barrydale and Swartberg (Table 1). In addition, a marginally significant negative quadratic coefficient was also observed at Swartberg, suggestive of stabilizing selection (Table 1). However, a lack of a clear global optimum from the thin plate spline, suggests that perhaps this could be a saturating effect, where floral tubes beyond a certain length do not confer an extra pollen receipt advantage (Fig. 4D, Fig. 3).

At Barrydale, pollen receipt surfaces differed significantly depending on whether the flowers were visited by bees or long proboscid flies (Barrydale: χ^2^ =17.65, edf =1.00, p <0.001). We only found evidence for positive linear selection when flowers at Barrydale were visited by long proboscid flies (Table 1). These results suggest that bees transfer pollen more evenly across flowers with different tube lengths, but that pollen deposition by flies strongly increases with tube length. In particular, the pollen receipt surfaces suggest that pollen receipt starts to increase very quickly as floral tubes approach 50mm in length (Fig. 4E).

At Swartberg, pollen receipt surfaces also differed significantly depending on whether the flowers were visited by bees or long proboscid flies (Swartberg: χ^2^ =3.36, edf =0.89, *P* =0.033, Fig. 4F). In particular, bees appeared to deposit much less pollen than flies across all phenotypes and have a much flatter pollen receipt surface than the one generated by flies. Nevertheless, there was evidence for a significant stabilizing selection coefficient associated with bee visits, irrespective of whether an outlier was excluded or not. Furthermore, a marginally significant linear relationship for bees was rendered non-significant after a single outlier was excluded (Table 1). Long proboscid fly pollinators appeared to be associated with both linear and stabilizing components of selection on tube length (Table 1). However, stabilizing selection may actually be indicative of a saturating effect predicted in Fig. 3 and observed in Fig. 4F..

## Discussion

Pollen receipt surfaces of *N. humilis* populations visited by bees versus flies were highly divergent (Fig. 4A and B). The short functional body lengths of bees facilitated greater pollen deposition onto short-styled flowers than long-styled flowers in Swellendam. This is intuitive as shorter styles are likely to make better contact with short bee bodies than longer styles (Fig. 3). In contrast, the more elongated anatomy of long proboscid flies in the Baviaanskloof increased pollen deposition on flowers with intermediate styles: short styles are unable to reach the bodies of foraging long proboscid flies whereas extremely long styles project past the bodies of foraging long proboscid flies (Fig, S2). The strong statistical differences in the pollen receipt surfaces of the bee versus the long proboscid fly populations, support the idea that functionally different pollinators impose divergent selection pressures on floral traits such as style length (Benitez-Vieyra et al. 2009). Indeed, Newman *et al*. (2015) demonstrated that divergent style lengths in *N. humilis* populations were locally adapted to the body dimensions of the dominant pollinators in those populations.

Similarly, for *T. revoluta*, the striking differences in bee and long-proboscid fly morphology were associated with different patterns of pollen receipt. Bees in both populations typically transferred fewer pollen grains per visit and the amount of pollen delivered did not differ as much in response to floral trait variation, resulting in flatter pollen receipt surfaces across floral tube lengths. As bee proboscis length was always shorter than the tube lengths of *T. revoluta*, bees likely pick up and deposit pollen more or less equally across all floral tube lengths when they push past the reproductive parts of *T. revoluta* in an attempt to extract nectar from the long tubes. Weak structure in these pollen receipt surfaces suggests that although bees can deposit pollen and play an important role in the pollination of *T. revoluta* (de Merxem et al 2009), their influence on tube length evolution is likely to be small.

This contrasts with long-proboscid flies visiting the same populations. Here, the highly structured pollen receipt surfaces suggest that long proboscid flies potentially exert strong selection on floral tube length through mechanical fit. The overall similarity in shape between global receipt surfaces (including bees) and the receipt surfaces generated by long proboscid flies (particularly in Barrydale but also Swartberg) further suggests the importance of long proboscid flies (but less-so bees) as potential agents of selection on floral tube length. In the Swartberg population, it was clear that pollen receipt was maximized in flowers with floral tubes that exceeded the visiting flies’ proboscis lengths and that beyond the average proboscis length of the flies, additional increases in tube length result in very little increase in pollen receipt. This saturation effect is intuitive because increasing the length of the floral tube beyond the length of the pollinator proboscis is not likely to increase the efficiency of pollen transfer, as all flies with proboscides shorter than floral tubes have to pass the stigma of *T. revoluta* to access the nectar (see predictions, Fig. 3).

Despite some general similarities in the pollen receipt surfaces imposed by long proboscid flies at Swartberg and Barrydale, there were also some important contrasts. These differences could primarily be explained by the three-fold difference in the proboscis lengths of the nemestrinid flies from these two communities. For example, the point at which the pollen receipt surface started to incline appeared to differ between the populations. When tubes were ca. 20 mm shorter than proboscides, flowers received virtually no pollen in both populations, but because fly proboscides differed across populations, this mismatch occurred at tube lengths of approximately 15 mm in the Swartberg and 50 mm at Barrydale.

Furthermore, in contrast to the Swartberg population, pollen receipt at the Barrydale population did not saturate but continued to increase with increasing floral tube length. We suggest that this is because it is easy to detect saturation effects when tube lengths are much longer than pollinator proboscis lengths as in the Swartberg population. But in the Barrydale population, we were not able to introduce plants with corolla tubes that were much longer than the fly proboscis lengths and so this saturation effect may have been harder to detect. These results suggest that floral morphology may evolve to enhance the mechanical fit between plant morphology and specific pollinators’ morphology [*sensu* Newman et al. (2015)].

### Geographic mosaics of pollinator selection

This manuscript isolates the potential for different kinds of pollinators to select on mechanical-fit traits within populations and across geographic pollinator mosaics. It helps us to visualize the contrasting effects of different pollinators and their differing potentials as drivers of selection on floral tube/style lengths within populations as well as in allopatric populations. For example, Anderson et al (2014) found that most *T. revoluta* populations are visited by long proboscid flies as well as bees. However, they found that floral tubes are much better matched to fly proboscis length at each population than to bee proboscis length. Our results, showing that pollen deposition by bees is both less effective and consistent across tube length variation, help us to visualize why floral tube length is apparently well adapted to long proboscid fly mouthparts rather than bee mouthparts. It also demonstrates how floral interactions with different functional pollinators can potentially generate coevolutionary hotspots between some species pairs but coldspots between others (Gomulkiewicz et al. 2000). In conjunction with pollination ecotype studies [e.g. Johnson et al. (2025)] and studies of pollinator selection in real time (Gervasi and Schiestl 2017), these results provide clarity of the mechanics behind selection driven by contrasting pollinator groups. Broadly, it also demonstrates how the contrasting effects of different pollinators may give rise to divergent selection on floral morphology when pollinators vary geographically as envisaged by Grant and Grant (1965), Stebbins (1970).

### Conclusions

Instead of assuming that flowers simply adapt to the most effective pollinator in a population, Aigner (2001) and Ohashi et al. (2021) suggest that floral adaptation may be better-studied by visualizing performance surfaces generated by different pollinators. Their work suggests that each pollinator species may contribute to selection on floral traits and that a wider set of adaptive outcomes are likely to exist, other than adaptation to the most effective pollinator. However, we know little about the real shapes of these surfaces and often surfaces of different pollinators are simply thought of as non-overlapping peaks with strong trade-offs that drive specialization to either one pollinator or another (Stebbins 1970). Our study suggests that this is the case for *N. humilis*, where bees and flies exert divergent patterns of selection, likely resulting in trade-offs in the use of these alternate pollinators and preventing the “effective” use of both pollinator groups within a population (Moir et al. 2025). Alternatively, overlapping performance surfaces may be important in facilitating shifts between different pollinator groups and consequent pollinator driven floral divergence. Here, we show in *T. revoluta* that performance surfaces of very different pollinators can be broadly overlapping, allowing both to contribute to plant reproductive fitness across a broad range of phenotypes. Importantly, the broad, flat shape of the bee performance surface allows flowers to adapt to fly morphology without reducing the contributions made by bees. Under normal circumstances, the elongation of floral tubes is expected to exclude pollinators with short proboscides [e.g., Haber and Frankie (1989)].

However, in this system, nectar wells up the long floral tubes of *T. revoluta*, facilitating the continued visitation by solitary bees and potentially contributing to seed set in years when long proboscid flies are rare or absent. The upwelling of nectar in this system can be viewed as a modifier trait [*sensu* Ohashi et al. (2021), Wang et al. (2023)], because it mitigates adaptive trade-offs of different pollinators allowing apparently specialized floral morphology to be effectively visited by very different pollinators. Importantly the flat performance surface of the bees suggests that floral tube length will always adapt to fly proboscis length, irrespective of how effective or numerous the bees are.

## Acknowledgements

We thank the Stellenbosch University Biodiversity and Ecology Honours classes of 2014 and 2015 for contributing to data collection of *Tritoniopsis revoluta*. Thanks to Michael Morrissey, Pascal Marrot, and Katharine Khoury for statistical input. We thank Cape Nature and the Eastern Cape Conservation board for collecting permits. We would like to thank Theo Busschau and Caitlin von Witt for their photographs of long proboscid flies in the field. We would also like to thank the National Research Foundation for the financial assistance given to BA over the course of this study (grants 87734, 105987, 137988) and to the Claude Leon Foundation (EN).

## Statement of authorship

All authors contributed to the generation of ideas, data collection, and interpretation of results. All data analysis was done by EN. EN wrote the first draft, with input from all authors thereafter.

## References

Aigner, P. A. 2001. Optimality modeling and fitness trade-offs: when should plants become pollinator specialists? Oikos 95:177–184.

Anderson, B. and S. D. Johnson. 2008. The geographical mosaic of coevolution in a plant–pollinator mutualism. Evolution 62:220–225.

Anderson, B., P. Ros, T. Wiese, and A. Ellis. 2014. Intraspecific divergence and convergence of floral tube length in specialized pollination interactions. Proceedings of the Royal Society B: Biological Sciences 281:20141420.

Arnold, S. J. 2003. Performance surfaces and adaptive landscapes. Integrative and comparative biology 43:367–375.

Benitez-Vieyra, S., A. Medina, and A. A. Cocucci. 2009. Variable selection patterns on the labellum shape of Geoblasta pennicillata, a sexually deceptive orchid. Journal of Evolutionary Biology 22:2354–2362.

de Merxem, D. G., B. Borremans, M. D. Jäger, T. Johnson, M. Jooste, P. Ros, R. Zenni, A. Ellis, and B. Anderson. 2009. The importance of flower visitors not predicted by floral syndromes. South African Journal of Botany 75:660–667.

Gervasi, D. D. and F. P. Schiestl. 2017. Real-time divergent evolution in plants driven by pollinators. Nature communications 8:14691.

Gomulkiewicz, R., J. N. Thompson, R. D. Holt, S. L. Nuismer, and M. E. Hochberg. 2000. Hot spots, cold spots, and the geographic mosaic theory of coevolution. The American Naturalist 156:156–174.

Grant, V. and K. A. Grant. 1965. Flower pollination in the Phlox family.

Haber, W. and G. Frankie. 1989. A tropical hawkmoth community: Costa Rican dry forest Sphingidae. Biotropica:155–172.

Hartig, F. and M. F. Hartig. 2017. Package ‘dharma’. R package.

Johnson, S. D. 2025. Pollination ecotypes and the origin of plant species. Proceedings B 292:20242787.

Johnson, S. D., T. van der Niet, E. Newman, N. Hobbhahn, and B. Anderson. 2025. Geographical variation in flower colour of a food-deceptive orchid reflects local pollinator preferences. Annals of Botany:mcaf074.

Lande, R. and S. J. Arnold. 1983. The measurement of selection on correlated characters. Evolution:1210-1226.

Mitchell-Olds, T. and R. G. Shaw. 1987. Regression analysis of natural selection: statistical inference and biological interpretation. Evolution 41:1149–1161.

Moir, M., H. Butler, C. Peter, T. Dold, and E. Newman. 2025. A test of the Grant-Stebbins pollinator shift model of floral evolution. New Phytologist In Press.

Morrissey, M. B. and I. B. Goudie. 2022. Analytical results for directional and quadratic selection gradients for log-linear models of fitness functions. Evolution 76:1378–1390.

Muchhala, N. and J. D. Thomson. 2009. Going to great lengths: selection for long corolla tubes in an extremely specialized bat–flower mutualism. Proceedings of the Royal Society B: Biological Sciences 276:2147–2152.

Newman, E., J. Manning, and B. Anderson. 2014. Matching floral and pollinator traits through guild convergence and pollinator ecotype formation. Annals of Botany 113:373–384.

Newman, E., J. Manning, and B. Anderson. 2015. Local adaptation: Mechanical fit between floral ecotypes of Nerine humilis (Amaryllidaceae) and pollinator communities. Evolution 69:2262–2275.

Ohashi, K., A. Jürgens, and J. D. Thomson. 2021. Trade-off mitigation: a conceptual framework for understanding floral adaptation in multispecies interactions. Biological Reviews.

Pauw, A., J. Stofberg, and R. J. Waterman. 2009. Flies and flowers in Darwins race. Evolution 63:268–279.

Ripley, B., B. Venables, D. M. Bates, K. Hornik, A. Gebhardt, D. Firth, and M. B. Ripley. 2013. Package ‘mass’. Cran r 538:113–120.

Robertson, J. L. and R. Wyatt. 1990. Evidence for pollination ecotypes in the yellow-fringed orchid, Platanthera ciliaris. Evolution 44:121–133.

Ros, P., A. G. Ellis, and B. Anderson. 2011. Maintenance of sympatric floral tube-length variation in a Cape irid. Biological journal of the Linnean Society 104:129–137.

Schluter, D. and D. Nychka. 1994. Exploring fitness surfaces. The American Naturalist 143:597–616.

Stebbins, G. L. 1970. Adaptive radiation of reproductive characteristics in angiosperms, I: pollination mechanisms. Annual review of ecology and systematics:307-326.

Sun, M., K. Gross, and F. P. Schiestl. 2014. Floral adaptation to local pollinator guilds in a terrestrial orchid. Annals of botany 113:289–300.

Trunschke, J., N. Sletvold, and J. Ågren. 2020. Manipulation of trait expression and pollination regime reveals the adaptive significance of spur length. Evolution 74:597–609.

Van der Niet, T., R. Peakall, and S. D. Johnson. 2014. Pollinator-driven ecological speciation in plants: new evidence and future perspectives. Annals of Botany 113:199–212.

Wade, M. J. and S. Kalisz. 1990. The causes of natural selection. Evolution 44:1947–1955.

Wang, B., Z.-Y. Tong, Y.-Z. Xiong, X.-F. Wang, W. Scott Armbruster, and S.-Q. Huang. 2023. The evolution of flower–pollinator trait matching, and why do some alpine gingers appear to be mismatched? Annals of Botany 132:1073–1088.

Waterman, R., H. Sahli, V. A. Koelling, K. Karoly, and J. K. Conner. 2023. Strong evidence for positive and negative correlational selection revealed by recreating ancestral variation. Evolution 77:264–275.

Wood, S. N. 2013. On p-values for smooth components of an extended generalized additive model. Biometrika 100:221–228.

Wood, S. N. 2017. Generalized additive models: an introduction with R. CRC press.

